# Repeat and haplotype aware error correction in nanopore sequencing reads with DeChat

**DOI:** 10.1101/2024.05.09.593079

**Authors:** Yichen Li, Enlian Chen, Jialu Xu, Wenhai Zhang, Xiangxiang Zeng, Yuansheng Liu, Xiao Luo

## Abstract

Error self-correction is a pivotal first step in the analysis of long-read sequencing data. However, most existing methods for this purpose are primarily tailored for noisy sequencing data with error rates exceeding 5%, often collapsing true variants in repeats and haplotypes. Alternatively, some methods are heavily optimized for PacBio HiFi reads, leaving a gap in methods specifically designed for Nanopore R10 reads basecalled with high accuracy or super accuracy models, which typically have error rates below 2%. Here, we introduce DeChat, a novel approach specifically designed for Nanopore R10 reads. DeChat enables repeat- and haplotype-aware error correction, leveraging the strengths of both de Bruijn graphs and variant-aware multiple sequence alignment to create a synergistic approach. This approach avoids read overcorrection, ensuring that variants in repeats and haplotypes are preserved while sequencing errors are accurately corrected. Benchmarking experiments reveal that reads corrected using DeChat exhibit substantially fewer errors, ranging from several times to two orders of magnitude lower, compared to the current state-of-the-art approaches. Furthermore, the application of DeChat for error correction significantly improves genome assembly across various aspects. DeChat is implemented as a highly efficient, standalone, and user-friendly software and is publicly available at https://github.com/LuoGroup2023/DeChat.

## Introduction

In recent years, third-generation sequencing (TGS) technologies, such as single-molecule real-time (SMRT) sequencing by Pacific Biosciences (PacBio) and nanopore sequencing by Oxford Nanopore Technologies (ONT), also known as long-read sequencing, have emerged. These technologies offer significant advantages in terms of read length (1) and PCR bias compared to Next Generation Sequencing (NGS). Consequently, they have greatly improved various applications in genomics, including genome assembly (2; 3; 4; 5; 6), variant calling (7; 8; 9), haplotype phasing (10), and taxonomic profiling (11). While long reads initially suffered from significantly high sequencing error rates ranging from 5% to 15% (1), their sequencing technologies have undergone rapid evolution. Recently, PacBio has been able to generate highly accurate (99.8%) long high-fidelity (HiFi) reads (12), and ONT R10 reads basecalled with high accuracy (HAC) or super accuracy (SUP) models already achieve over 98% modal raw read accuracy (13; 14). Remarkably, ONT R10.4 duplex reads basecalled with the SUP model can achieve comparable accuracy to HiFi reads, albeit at the cost of a dramatic reduction in sequencing throughput. Error self-correction is typically the initial and crucial step in the analysis of long-read sequencing data, particularly in genome assembly. It has become a standard procedure in haplotype-aware genome assembly (15; 16; 17; 18; 19; 20; 21) and strain-aware metagenome assembly (22; 23; 24).

However, most existing long-read error self-correction methods are tailored specifically for noisy sequencing data with error rates exceeding 5% (15; 25; 26; 27; 28; 29). These tools typically generate corrected long reads by computing a consensus sequence for each read, which can result in the loss of genetic variations across different repeats, haplotypes, or strains. While our recent method, VeChat (30), aims to address this issue by utilizing variation graphs, it suffers from poor speed primarily due to the inefficiency of variation graph construction. Although PacBio HiFi reads are relatively straightforward to correct due to their high accuracy, the corresponding error correction methods are often integrated into genome assemblers(16; 17; 18; 22; 23). Furthermore, these methods are heavily tailored for HiFi reads and are not directly applicable to nanopore sequencing data, as the two data types exhibit different error rates and profiles. Consequently, there is a paucity of methods specifically designed for ONT R10 reads (with error rates below 2%) that can simultaneously achieve high performance in terms of accuracy and efficiency.

To address this gap, we have developed a self-correction method called DeChat ([D]e Bruijn graph-based [e]rror [C]orrection in [ha]plo[t]ypes). DeChat corrects sequencing errors in ONT R10 long reads in a manner that is aware of repeats, haplotypes or strains. Methodologically, it combines the concepts of de Bruijn graphs (dBG) and variant-aware multiple sequence alignment (MSA), leveraging the strengths of both strategies to create a synergistic approach. We have evaluated DeChat and compared it with other prominent tools on various datasets with different settings, including haploid genomes, polyploid genomes, and metagenomes. Benchmarking experiments have demonstrated that our approach, DeChat, essentially achieves the optimal performance in terms of error rate on both simulated and real data. Furthermore, the utilization of DeChat for error correction considerably improves (meta)genome assembly across various aspects.

## Results

### Approach

We have designed and implemented DeChat, a highly efficient method for repeat- and haplotype-aware error self-correction, tailored specifically for ONT R10 long reads. The core principle of DeChat lies in the synergistic integration of de Bruijn graphs and variant-aware multiple sequence alignment (MSA). On one hand, de Bruijn graphs facilitate efficient pre-correction of raw reads. On the other hand, variant-aware MSA enables repeat- or haplotype-specific consensus generation, effectively preventing overcorrection of variations among different repeats or haplotypes in samples of higher, known or unknown ploidy.

Figure 1 illustrates the workflow of DeChat, which comprises two distinct stages. In the first stage, DeChat divides raw reads into small kmers and eliminates those with extremely low frequencies (indicating potential errors). Subsequently, it constructs a compacted de Bruijn graph (dBG). This approach is justified by the fact that the error rate of ONT R10 long reads typically remains below 2%. Each raw read is then aligned to the compacted dBG to identify the optimal alignment path, resulting in the pre-corrected read (as depicted in the left panel of Figure 1).

**Figure 1.**
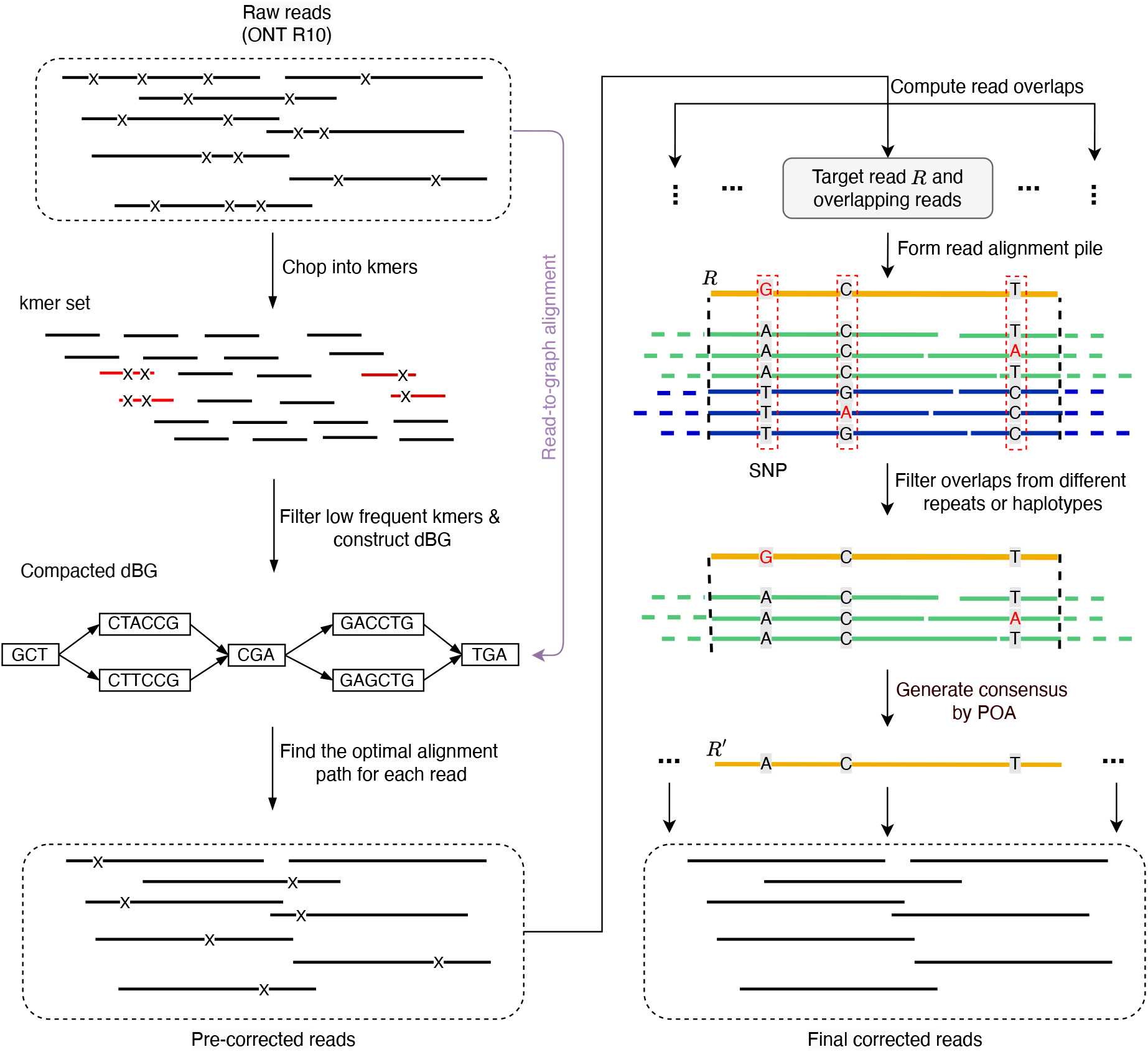
Workflow of DeChat. The left and the right panel indicates the first and the second stage, respectively. The forks in reads represent sequencing errors. In the right panel, the orange line indicates the target read, and the green and blue reads are from different repeats or haplotypes. POA: partial order alignment.

The second stage (depicted in the right panel) begins with computing all-vs-all overlaps among the pre-corrected reads. A target read *R* (to be corrected) is selected, and its overlapping reads are gathered to form a read alignment pile. DeChat identifies heterozygous sites along the target read and retains only the overlapping reads that share the same alleles as the target read. This step ensures the purity of reads in the alignment pile, ensuring that they originate from the same repetitive region or haplotype. This is crucial to prevent overcorrection. Subsequently, DeChat employs a multiple sequence alignment approach (via partial order alignment algorithm) to generate a consensus sequence, serving as the corrected target read *R*^*′*^. Each pre-corrected read undergoes this correction process in the same manner. Notably, to further enhance error correction, the second stage can be iterated several times (default 3) to produce the final corrected reads.

### Benchmarking results

Figure 2A presents the benchmarking results for error correction in simulated and real ONT reads derived from genomes with varying ploidies, specifically 2, 3, and 4. DeChat demonstrates a significant reduction in error rates, achieving approximately 10 ∼ 236, 3.5 ∼ 5.8 and 2 times lower error rates on pseudo polyploid (*E. coli*), potato, and human (HG002) genomes, respectively. Notably, it maintains superior or comparable performance across other metrics, including haplotype coverage, the number of corrected reads and misassemblies, and the length of corrected reads (as evidenced by N50/NGA50). Detailed information can be found in Supplementary Tables 1 and 2.

**Figure 2.**
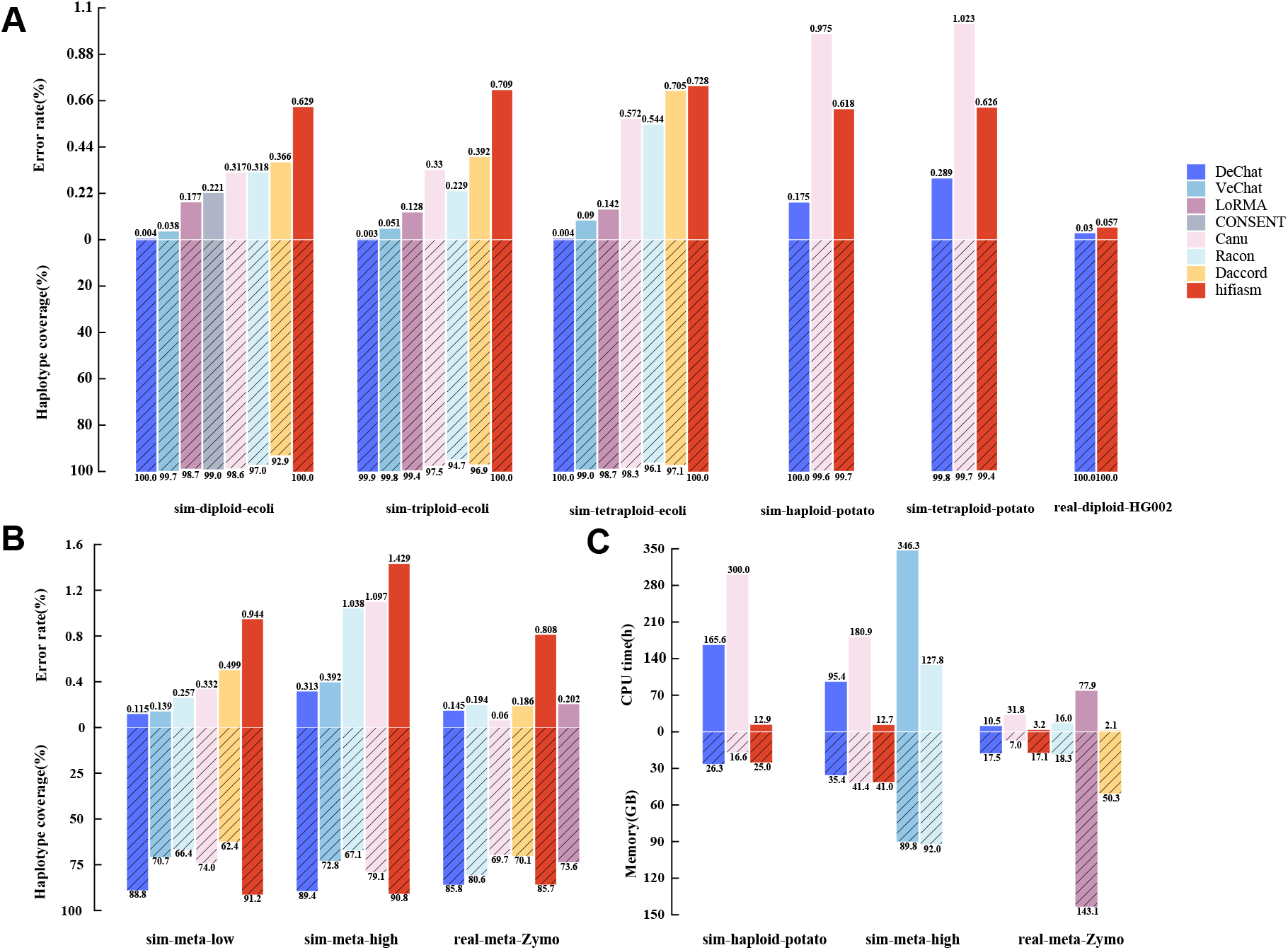
Benchmarking results of error correction. (A) Error correction results of haploid and polyploid genomes. (B) Error correction results of metagenomes. (C) Running time and peak memory usages of error correction tools. Notably, some tools were unable to process certain datasets due to either failure to execute them or the inability to complete the process within an acceptable time frame. The results of these tools are therefore not shown for those specific datasets.

Figure 2B displays the benchmarking results for error correction in simulated and real ONT reads from metagenomes with varying degrees of complexity. DeChat achieves notable improvements in error rates, reducing them by approximately 1.2 ∼ 8, 1.3 ∼ 5 and 1.3 ∼ 5.5 (except Canu) times on low-complexity metagenome, high-complexity metagenome, and Zymo (mock community) datasets, respectively. Simultaneously, it maintains comparable performance across other aspects (see Supplementary Table 3 for detailed information). Notably, Canu performs slightly better than DeChat on real-meta-Zymo dataset in terms of error rate. However, it is at the significant cost of a 16% reduction in haplotype coverage.

DeChat exhibits slower performance compared to hifiasm but generally outperforms all other error correction tools, as evident in Figure 2C. Notably, our experiments highlight that only DeChat, hifiasm, and Canu are capable of handling big datasets from large genomes, such as potato and human genomes. Nevertheless, DeChat offers a notable advantage in terms of speed, achieving approximately twice the speed of Canu’s error correction on the sim-haploid-potato dataset, as shown in Figure 2C. Furthermore, DeChat’s peak memory usage is comparable to that of hifiasm, making it a viable option for resource-constrained environments.

### Improving genome assembly

To investigate how correcting errors in reads influences the quality of de novo genome assemblies, we conducted a series of experiments using hifiasm and hifiasm-meta, two leading assemblers for polyploid genomes and metagenomes, respectively. We ran these assemblers on datasets described earlier, both with and without first correcting the reads using DeChat. The results are unequivocal: hifiasm and hifiasm-meta significantly benefit from the use of DeChat-corrected reads (see Figure 3). Across the vast majority of datasets, we observe improved haplotype coverage, longer contigs (measured by N50/NGA50), lower error rates, and fewer misassemblies when DeChat is applied prior to assembly. See Supplementary Tables 4 ∼ 7 for the details.

**Figure 3.**
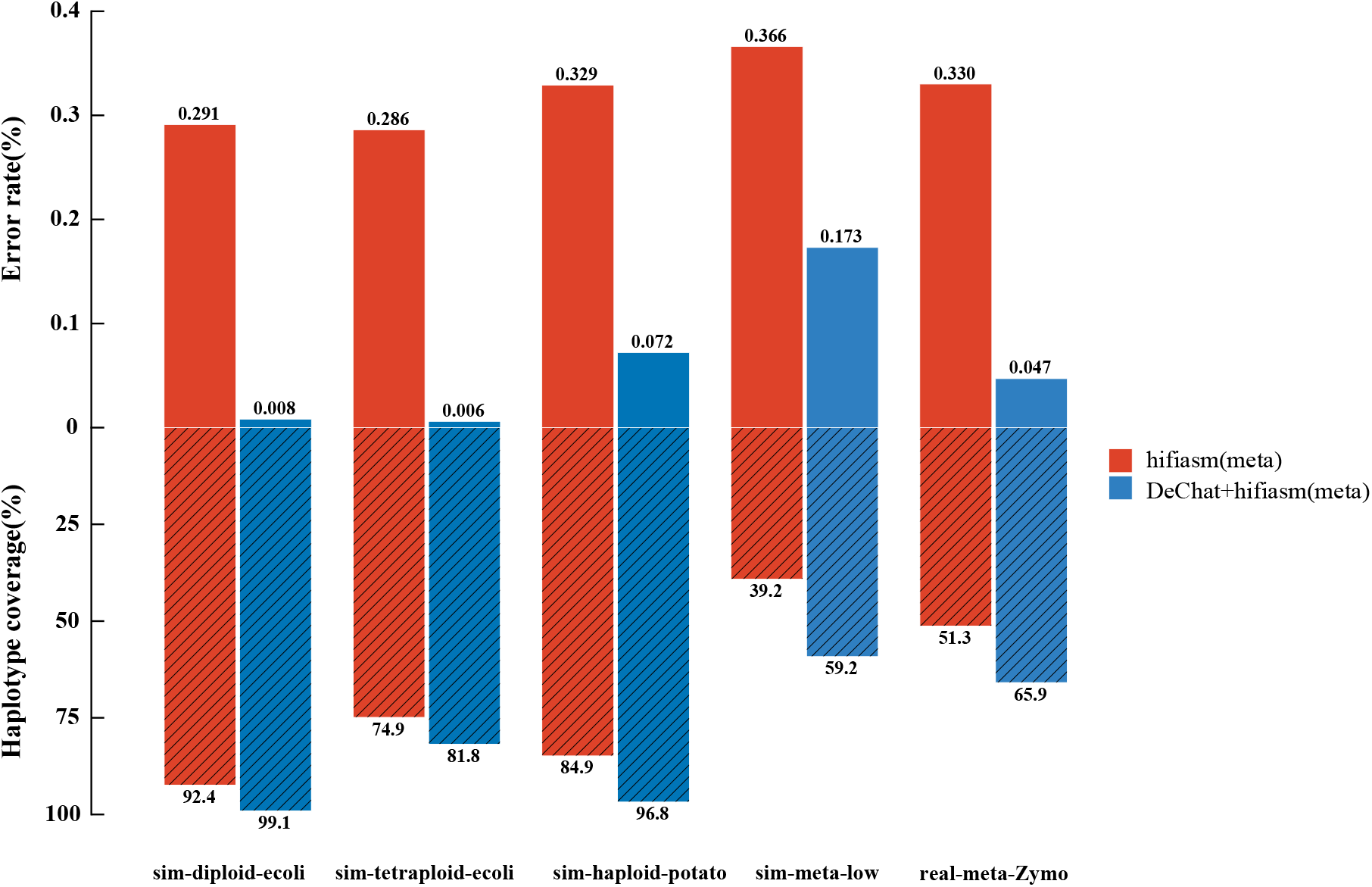
Assembly results of either using DeChat for error correction or not. Note that we used hifiasm for haploid or polyploid genome assembly, and hifiasm-meta for metagenome aseembly, respectively.

## Discussion

We have introduced DeChat, a novel approach that performs repeat- and haplotype-aware error correction specifically designed for nanopore sequencing long reads. To the best of our knowledge, DeChat stands as the first method tailored and optimized for ONT R10 reads basecalled using the HAC or SUP models, which boast an error rate below 2%. This advancement is particularly timely and relevant in the current era, where sequencing accuracy and cost-effectiveness of ONT platforms are constantly improving.

The key methodological advancements of DeChat lie in its judicious integration of the efficiency of de Bruijn graphs and the sensitivity of variant-aware multiple sequence alignment. This innovative combination not only addresses the computational bottlenecks encountered by traditional approaches but also effectively mitigates the risk of overcorrection, a common pitfall in error correction algorithms. By harnessing these strengths, DeChat offers a robust and efficient solution for enhancing the quality of nanopore sequencing data, paving the way for more accurate and reliable downstream analyses.

Our findings strongly indicate the superiority of DeChat in correcting errors in nanopore sequencing long reads. Across most of benchmarking scenarios, DeChat significantly reduces error rates by a factor of several times to two orders of magnitude compared to leading competitive methods. This remarkable improvement in error correction is achieved without compromising the haplotype completeness of the reads. In fact, DeChat is particularly adept at preventing overcorrection of variations between distinct repeats or haplotypes, thereby preserving the haplotype identity of the reads.

Moreover, our study demonstrates that the use of DeChat for read correction prior to de novo genome assembly significantly enhances the quality of the resulting assemblies. This is unsurprising given that DeChat effectively distinguishes haplotype-specific variations from sequencing errors, thus maintaining haplotype information in the corrected reads. Notably, the application of DeChat-corrected reads enables the successful utilization of assembly tools like hifiasm, which crucially rely on high-quality input data. This underscores the significance and utility of DeChat in facilitating accurate and haplotype-aware genome assemblies.

However, while the corrected reads by DeChat have improved the assembly process, it is evident that further advancements are still required in the development of haplotype-aware assemblers, especially when dealing with complex heterogeneous genomes like polyploid genomes and metagenomes. These challenges represent promising areas for future exploration, but they lie beyond the scope of this work. Undoubtedly, there is ample room for future improvements. Notably, the first stage of DeChat consumes a substantial amount of time, necessitating further acceleration. Recent studies, including (17; 23), demonstrate that the utilization of minimizer-based sparse de Bruijn graphs significantly enhances the efficiency of genome assembly from accurate long reads. Consequently, exploring the potential of sparse de Bruijn graphs within the first stage of DeChat holds considerable interest for future endeavors.

## Methods

### Implementation

We have developed DeChat as a highly efficient, stand-alone, and user-friendly software. The core of DeChat is built upon several established methods, each specializing in a specific aspect of the error correction process. In the first stage, DeChat rapidly constructs a compacted de Bruijn graph (dBG) from raw reads in a parallel manner, leveraging a minimizer hashing strategy (31). Subsequently, it aligns the raw reads to the compacted dBG and identifies the optimal alignment paths, which spell out the pre-corrected reads. DeChat implements the read-to-graph alignment process by employing an efficient depth-first search algorithm (32). In the second stage, DeChat performs variant-aware multiple sequence alignment by adapting the error correction module from hifiasm (18) to generate the final corrected reads. By integrating and optimizing these components from diverse approaches, we have packaged them into a seamless, standalone software that is straightforward to install and use. DeChat provides a comprehensive solution for error correction in nanopore long-read sequencing data, tailored to meet the diverse needs of a wide range of users and applications.

### Datasets

We have constructed pseudo diploid, triploid, and tetraploid genomes by mixing strains of *Escherichia coli* (*E. coli*) bacteria. These genomes are designated as “sim-diploid-ecoli”, “sim-triploid-ecoli”, and “sim-tetraploid-ecoli”, respectively. Additionally, we utilized a high-quality tetraploid potato genome from a recent study (33) to construct a real tetraploid and a real haploid plant genome, named “sim-tetraploid-potato” and “sim-haploid-potato”, respectively. It is noteworthy that while the genomes in potato datasets are authentic, the ONT R10 reads are synthetic.

Furthermore, we employed CAMISIM (34) to simulate two metagenomic datasets, each with distinct complexities. The low complexity dataset, named “sim-meta-low”, comprises 30 species (60 strains) with strain relative abundances ranging from 0.30% to 6.43%. Whereas the high complexity dataset, named “sim-meta-high”, includes 373 species (1000 strains) with strain relative abundances ranging from 0.04% to 0.30%. The microbial genomes involved in both datasets were sourced from complete genome sequences deposited in RefSeq.

For read simulation, we used PBSIM2 (35), a widely used tool, in its sampling-based simulation mode for all synthetic datasets. The read template file was derived from a recent ONT long-read genome sequencing study (14), which reported an error rate of approximately 2%.

In addition, we downloaded the real ONT R10.4.1 duplex reads of the human genome (HG002) from https://s3-us-west-2.amazonaws.com/human-pangenomics/index.html?prefix=submissions/0CB931D5-AE0C-41HG002_R1041_Duplex_Dorado/Dorado_v0.1.1/stereo_duplex/ and the real ONT R10.4.1 reads of a metagenome (Zymo) from this study(14). These datasets are designated as “real-diploid-HG002” and “real-meta-Zymo”, respectively.

### Evaluation

To ensure a fair and accurate comparison, we deliberately selected various representative methods for long-read error self-correction, including the error correction module of Canu (15), Racon (25), LoRMA (26), CONSENT (28), Daccord (29) and VeChat (30). Additionally, we included the error correction module of hifiasm (18) in our comparison, as it is a widely utilized haplotype-aware HiFi assembler and has demonstrated superior performance in terms of error correction (36).

For evaluating the performance of long-read error correction and genome assembly, we utilized QUAST (37). The primary metrics we report are error rate (ER) and haplotype coverage (HC). The error rate is calculated as the sum of mismatch rate and indel rate when aligning the corrected reads or contigs to the reference genomes. Haplotype coverage, on the other hand, represents the percentage of aligned bases in the ground truth haplotypes that are covered by the corrected reads or contigs. For further details regarding the evaluation process, please refer to the Supplementary materials.

## Data availability

The genomes and simulated sequencing reads generated by this study can be downloaded from Zenodo DOI: 10.5281/zenodo.10995721 (https://doi.org/10.5281/zenodo.10995722).

## Code availability

The source code of DeChat is GPL-3.0 licensed, and publicly available at https://github.com/LuoGroup2023/DeChat.

## Acknowledgements

This study was supported by the Natural Science Foundation of Hunan Province (Grant No. 2024JJ4008); Fundamental Research Funds for the Central Universities (Grant No. 541109030062); the National Natural Science Foundation of China (Grant No. 62372159, 62102140, 62172002); the Science and Technology Innovation Program of Hunan Province (2022RC1100).

## Author contributions

Xiao Luo conceived this study and designed the method. Yichen Li implemented the software. Xiao Luo, Yuansheng Liu and Xiangxiang Zeng wrote and revised the manuscript. Yichen Li, Enlian Chen, Jialu Xu, Wenhai Zhang conducted the data analysis. All authors read and approved the final version of the manuscript.

