## Supplementary Material for "Repeat and haplotype aware error correction in nanopore sequencing reads with DeChat"

#### Supplementary Tables and Figures

| Method | #Reads | Haplotype coverage (%) | N50 (bp) | NGA50 (bp) | Error rate (%) | Mismatch (%) | Indel (%) | # Misassemblies |
|---|---|---|---|---|---|---|---|---|
| sim-diploid-ecoli | | | | | | | | |
| DeChat | 21019 | 100.0 | 8215 | 45553 | 0.004 | 0.002 | 0.002 | 0 |
| VeChat | 16426 | 99.7 | 8987 | 45251 | 0.038 | 0.033 | 0.004 | 1 |
| LoRMA | 20472 | 98.7 | 7282 | 29944 | 0.177 | 0.089 | 0.088 | 25 |
| CONSENT | 21003 | 99.0 | 8239 | 45159 | 0.221 | 0.206 | 0.015 | 68 |
| Canu | 19834 | 98.6 | 8381 | 45020 | 0.317 | 0.297 | 0.020 | 14 |
| Racon | 20747 | 97.0 | 8283 | 45522 | 0.318 | 0.285 | 0.033 | 42 |
| Daccord | 24298 | 92.9 | 6867 | 9891 | 0.366 | 0.352 | 0.013 | 129 |
| hifiasm | 21019 | 100.0 | 8216 | 45471 | 0.629 | 0.170 | 0.459 | 0 |
| Raw | 21019 | 100.0 | 8230 | 45537 | 1.821 | 0.439 | 1.382 | 0 |
| sim-triploid-ecoli | | | | | | | | |
| DeChat | 28505 | 99.9 | 8956 | 47233 | 0.003 | 0.002 | 0.002 | 0 |
| VeChat | 21782 | 99.8 | 9963 | 46892 | 0.051 | 0.047 | 0.004 | 12 |
| LoRMA | 28389 | 99.4 | 7977 | 29930 | 0.128 | 0.061 | 0.067 | 15 |
| Racon | 27914 | 94.7 | 9058 | 46925 | 0.229 | 0.208 | 0.021 | 204 |
| Canu | 26453 | 97.5 | 9211 | 46498 | 0.330 | 0.312 | 0.018 | 47 |
| Daccord | 28531 | 96.9 | 8906 | 46669 | 0.392 | 0.380 | 0.012 | 193 |
| hifiasm | 28505 | 100.0 | 8967 | 47070 | 0.709 | 0.186 | 0.522 | 0 |
| Raw | 28505 | 100.0 | 8980 | 47231 | 1.892 | 0.455 | 1.437 | 0 |
| sim-tetraploid-ecoli | | | | | | | | |
| DeChat | 38866 | 100.0 | 8965 | 47092 | 0.004 | 0.002 | 0.003 | 0 |
| VeChat | 29815 | 99.0 | 9957 | 46755 | 0.090 | 0.083 | 0.007 | 10 |
| LoRMA | 38485 | 98.7 | 7856 | 29935 | 0.142 | 0.084 | 0.058 | 36 |
| Racon | 37934 | 96.1 | 9058 | 46459 | 0.544 | 0.500 | 0.044 | 216 |
| Canu | 33108 | 98.3 | 9592 | 46446 | 0.572 | 0.542 | 0.030 | 61 |
| Daccord | 38890 | 97.1 | 8937 | 46609 | 0.705 | 0.688 | 0.017 | 166 |
| hifiasm | 38866 | 100.0 | 8971 | 46997 | 0.728 | 0.196 | 0.532 | 0 |
| Raw | 38866 | 100.0 | 8995 | 47047 | 1.884 | 0.459 | 1.425 | 0 |

**Supplementary Table 1.** Error correction benchmarking results for simulated Ecoli genomes of various ploidy (ploidy=2,3,4). The average sequencing coverage per haplotype is 10x and sequencing error rate is approximately 2%. "#Reads" indicates the number of corrected reads. The error rate is equal to the sum of mismatch and indel rate. The results are sorted by the error rate in ascending order.

| Method | #Reads | Haplotype coverage (%) | N50 (bp) | NGA50 (bp) | Error rate (%) | Mismatch (%) | Indel (%) | # Mis-assem-blies |
|---|---|---|---|---|---|---|---|---|
| sim-haploid-potato | | | | | | | | |
| DeChat | 1511276 | 100.0 | 8841 | 47953 | 0.175 | 0.066 | 0.109 | 132 |
| hifiasm | 1401815 | 99.7 | 8646 | 45884 | 0.618 | 0.162 | 0.456 | 19 |
| Canu | 1317101 | 99.6 | 8841 | 45701 | 0.975 | 0.612 | 0.363 | 2454 |
| Raw | 1511276 | 100.0 | 8872 | 47797 | 1.857 | 0.453 | 1.404 | 69 |
| sim-tetraploid-potato | | | | | | | | |
| DeChat | 6075084 | 99.8 | 8833 | 47924 | 0.289 | 0.096 | 0.193 | 854 |
| hifiasm | 5633198 | 99.4 | 8617 | 45751 | 0.626 | 0.167 | 0.460 | 158 |
| Canu | 5734712 | 99.7 | 9007 | 47427 | 1.023 | 0.722 | 0.300 | 21558 |
| Raw | 6075084 | 100.0 | 8862 | 47742 | 1.856 | 0.454 | 1.402 | 648 |
| real-diploid-HG002 | | | | | | | | |
| DeChat | 4287987 | 100.0 | 35350 | 67816 | 0.030 | 0.008 | 0.022 | 59129 |
| hifiasm | 4288001 | 100.0 | 35345 | 67794 | 0.057 | 0.017 | 0.040 | 58630 |
| Raw | 4287993 | 100.0 | 35335 | 67771 | 0.180 | 0.050 | 0.130 | 58664 |

**Supplementary Table 2.** Error correction benchmarking results for simulated ONT reads of haploid and tetraploid potato genomes, as well as diploid HG002. The average sequencing coverage per haplotype for potato is 10x with a sequencing error rate of approximately 2%, and for HG002 it is 20x with a sequencing error rate of approximately 0.2%. "#Reads" indicates the number of corrected reads. The error rate is equal to the sum of the mismatch and indel rates. The results are sorted by the error rate in ascending order.

| Method | #Reads | Haplotype coverage (%) | N50 (bp) | NGA50 (bp) | Error rate (%) | Mismatch (%) | Indel (%) | # Mis-assem-blies |
|---|---|---|---|---|---|---|---|---|
| sim-meta-low | | | | | | | | |
| DeChat | 251477 | 88.8 | 8834 | 40580 | 0.115 | 0.064 | 0.052 | 34 |
| VeChat | 183864 | 70.7 | 9985 | 39387 | 0.139 | 0.108 | 0.030 | 1291 |
| Racon | 245054 | 66.4 | 8946 | 39444 | 0.257 | 0.204 | 0.053 | 2349 |
| Canu | 250076 | 74.0 | 8834 | 39746 | 0.332 | 0.242 | 0.090 | 1410 |
| Daccord | 256873 | 62.4 | 8389 | 36331 | 0.499 | 0.441 | 0.058 | 8019 |
| hifiasm | 251479 | 91.2 | 8848 | 40210 | 0.944 | 0.250 | 0.693 | 2 |
| Raw | 251479 | 92.4 | 8865 | 40217 | 1.867 | 0.454 | 1.413 | 2 |
| sim-meta-high | | | | | | | | |
| DeChat | 2137832 | 89.4 | 8872 | 28863 | 0.313 | 0.150 | 0.163 | 193 |
| VeChat | 1546498 | 72.8 | 10573 | 26969 | 0.392 | 0.240 | 0.152 | 18528 |
| Racon | 2052849 | 67.1 | 9044 | 25115 | 1.038 | 0.837 | 0.201 | 38313 |
| Canu | 2115564 | 79.1 | 8880 | 27470 | 1.097 | 0.863 | 0.234 | 12701 |
| hifiasm | 2137875 | 90.8 | 8898 | 28608 | 1.429 | 0.369 | 1.061 | 19 |
| Raw | 2137875 | 92.3 | 8905 | 28633 | 1.867 | 0.456 | 1.412 | 17 |
| real-meta-Zymo | | | | | | | | |
| Canu | 150769 | 69.7 | 10250 | 52399 | 0.060 | 0.040 | 0.020 | 9400 |
| Dechat | 233614 | 85.8 | 8797 | 52893 | 0.145 | 0.080 | 0.065 | 120786 |
| Daccord | 277995 | 70.1 | 6390 | 40602 | 0.186 | 0.151 | 0.034 | 37923 |
| Racon | 222356 | 80.6 | 9022 | 51989 | 0.194 | 0.147 | 0.047 | 96747 |
| LoRMA | 194965 | 73.6 | 8721 | 30148 | 0.202 | 0.109 | 0.093 | 33686 |
| hifiasm | 233614 | 85.7 | 8777 | 52152 | 0.808 | 0.428 | 0.379 | 103818 |
| Raw | 233614 | 85.7 | 8760 | 52061 | 1.700 | 0.876 | 0.824 | 103684 |

**Supplementary Table 3.** Error correction benchmarking results for simulated ONT reads of sim-meta-low (60 strains) and sim-meta-high (1000 strains), as well as Zymo real metagenomic dataset. "sim-meta-low" consists of strains with relative abundances ranging from 0.30% to 6.43%. "sim-meta-high" consists of strains with relative abundances ranging from from 0.04% to 0.30%. Additionally, the sequencing error rate is approximately 2%. "#Reads" indicates the number of corrected reads. The error rate is equal to the sum of mismatch and indel rate. The results are sorted by the error rate in ascending order.

| Method | Assembler | #Reads | Haplotype coverage (%) | N50 (bp) | NGA50 (bp) | Error rate (%) | Mismatch (%) | Indel (%) | # Mis-assem-blies |
|---|---|---|---|---|---|---|---|---|---|
| sim-diploid-ecoli | | | | | | | | | |
| DeChat | hifiasm | 40 | 99.1 | 537398 | 537398 | 0.008 | 0.004 | 0.004 | 0 |
| Raw | hifiasm | 111 | 92.4 | 139149 | 127606 | 0.291 | 0.087 | 0.204 | 8 |
| sim-triploid-ecoli | | | | | | | | | |
| DeChat | hifiasm | 68 | 69.1 | 1946151 | 1610698 | 0.008 | 0.004 | 0.004 | 1 |
| Raw | hifiasm | 86 | 66.0 | 277865 | 128913 | 0.406 | 0.110 | 0.296 | 1 |
| sim-tetraploid-ecoli | | | | | | | | | |
| DeChat | hifiasm | 117 | 81.8 | 546201 | 546200 | 0.006 | 0.003 | 0.003 | 0 |
| Raw | hifiasm | 143 | 74.9 | 261373 | 194410 | 0.286 | 0.082 | 0.204 | 4 |

**Supplementary Table 4.** Assembly benchmarking results for Ecoli genomes of various ploidy (ploidy=2,3,4) using hifiasm. The average sequencing coverage per haplotype is 10x, and the sequencing error rate is approximately 2%. "#Reads" indicates the number of corrected reads. The error rate is equal to the sum of mismatch and indel rate. The results are sorted by the error rate in ascending order.

| Method | Assembler | #Reads | Haplotype coverage (%) | N50 (bp) | NGA50 (bp) | Error rate (%) | Mismatch (%) | Indel (%) | # Mis-assem-blies |
|---|---|---|---|---|---|---|---|---|---|
| sim-meta-low | | | | | | | | | |
| DeChat | hifiasmmeta | 1767 | 59.2 | 274359 | 71431 | 0.173 | 0.109 | 0.064 | 124 |
| Raw | hifiasmmeta | 1603 | 39.2 | 127206 | - | 0.366 | 0.108 | 0.258 | 64 |
| sim-meta-high | | | | | | | | | |
| DeChat | hifiasmmeta | 29792 | 37.6 | 72700 | - | 0.451 | 0.266 | 0.185 | 3155 |
| Raw | hifiasmmeta | 17517 | 17.9 | 53692 | - | 0.900 | 0.257 | 0.643 | 1084 |

**Supplementary Table 5.** Assembly benchmarking results for sim-meta-low (60 strains) and sim-meta-high (1000 strains) using hifiasmmeta. "#Reads" indicates the number of corrected reads. The error rate is equal to the sum of mismatch and indel rate. The results are sorted by the error rate in ascending order.

| Method | Assembler | #Reads | Haplotype coverage (%) | N50 (bp) | NGA50 (bp) | Error rate (%) | Mismatch (%) | Indel (%) | # Mis-assem-blies |
|---|---|---|---|---|---|---|---|---|---|
| sim-tetraploid-potato | | | | | | | | | |
| DeChat | hifiasm | 17283 | 73.4 | 417853 | 276851 | 0.098 | 0.038 | 0.060 | 2304 |
| Raw | hifiasm | 21307 | 63.8 | 238033 | 136318 | 0.230 | 0.065 | 0.165 | 1971 |
| sim-haploid-potato | | | | | | | | | |
| DeChat | hifiasm | 5528 | 96.8 | 401207 | 570351 | 0.072 | 0.031 | 0.042 | 332 |
| Raw | hifiasm | 9366 | 84.9 | 123936 | 123543 | 0.329 | 0.086 | 0.243 | 365 |

**Supplementary Table 6.** Assembly benchmarking results for simulated ONT reads of haploid and tetraploid potato genomes using hifiasm. The average sequencing coverage per haplotype for potato is 10x with a sequencing error rate of approximately 2%. "#Reads" indicates the number of corrected reads. The error rate is equal to the sum of mismatch and indel rate. The results are sorted by the error rate in ascending order.

| Method | Assembler | #Reads | Haplotype coverage (%) | N50 (bp) | NGA50 (bp) | Error rate (%) | Mismatch (%) | Indel (%) | # Mis-assem-blies |
|---|---|---|---|---|---|---|---|---|---|
| real-diploid-HG002 | | | | | | | | | |
| Raw | hifiasm | 1154 | 96.8 | 51020825 | 7815285 | 0.033 | 0.016 | 0.017 | 1191 |
| DeChat | hifiasm | 1107 | 96.0 | 52513888 | 8059652 | 0.035 | 0.017 | 0.018 | 1268 |
| real-meta-Zymo | | | | | | | | | |
| DeChat | hifiasmmeta | 1178 | 65.9 | 97312 | 104661 | 0.047 | 0.027 | 0.020 | 1770 |
| Raw | hifiasmmeta | 816 | 51.3 | 91651 | 43326 | 0.330 | 0.169 | 0.161 | 971 |

**Supplementary Table 7.** Assembly benchmarking results for real-diploid-HG002 and real-meta-Zymo using hifiasm and hifiasm-meta, respectively. "#Reads" indicates the number of corrected reads. The error rate is equal to the sum of mismatch and indel rate. The results are sorted by the error rate in ascending order.

### Supplementary Methods

#### Metrics for evaluation

The genome assembly performance was evaluated by means of several commonly used metrics, routinely reported by QUAST V5.1.0 (Mikheenko *et al.*, 2018), as a prominent assembly evaluation tool. See below for specific explanations. Since error-corrected long reads have very similar properties with assembled contigs and for the purpose of unified comparison, we also used QUAST for evaluating the performance of long-read error correction. Corrected reads and contigs with length less than 500bp were filtered before evaluation. In particular, we ran the `quast.py` program with the option `--unique-mapping` appropriately taking into account that our data sets reflect mixed samples (such as polyploid genome or metagenome).

**Haplotype coverage (HC).** Haplotype coverage is the percentage of aligned bases in the ground truth haplotypes covered by corrected reads or contigs, which is used to measure the completeness of the given sequence data.

**N50 and NGA50.** N50 is defined as the length for which the collection of all corrected reads/contigs of that length or longer covers at least half the given sequences. NGA50 is similar to N50 but can only be calculated when the reference genome is provided. NGA50 only considers the aligned blocks (after breaking reads/contigs at misassembly events and trimming all unaligned nucleotides), which is defined as the length for which the overall size of all aligned blocks of this length or longer equals at least half of the reference haplotypes. Both N50 and NGA50 are used to measure the length distribution of corrected reads and the contiguity of the assemblies.

**Error rate (ER).** The error rate is equal to the sum of mismatch rate and indel rate when mapping the obtained corrected reads or contigs to the reference haplotype sequences.

**Number of misassemblies (#MA).** The misassembly event in corrected reads or assemblies indicates that left and right flanking sequences align to the true haplotypes with a gap or overlap of more than 1kbp, or align to different strands, or even align to different haplotypes or strains. Here, we report the total number of misassemblies in the given sequence data.

#### Commands and versions of tools used for comparison

- PBSIM2
```
pbsim --prefix $prefix --depth $depth --sample-fastq $fq_sample $reference --length-min 1000
```

- DeChat
```
dechat -i $raw_read -t $threads -o $corrected_read

#(real-meta-Zymo)
dechat -i $raw_read -t $threads -o -r1 3 -r 6 -e 0.04
```

- VeChat v1.1.0
```
vechat -o $corrected_read --platform ont $raw_read
```

- LoRMA v0.5
```
#LoRMA can only process reads shorter than 40,000 bp, therefore long reads need to be trimmed

LoRMA -reads $raw_read -output $corrected_read -discarded $discarded_read -nb-cores $threads \
-k $kmer
```

- Racon v1.5.0

```
minimap2 -x ava-ont --dual=yes $raw_read $raw_read -t $threads > $overlap.tmp.paf
awk '$11>=500' $overlap.tmp.paf > $overlap.tmp.filter.paf
fpa drop --same-name --internalmatch $overlap.tmp.filter.paf > $overlap
racon -f -t $threads $raw_read $overlap $raw_read > $tmp.fa
Filter out reads shorter than 1000bp from $temp.fa to obtain $correct_read
```

- hifiasm v0.19.8-r603

```
hifiasm -o $corrected_read -t 32 $raw_read
```

- Canu v2.2

```
canu -p $prefix -d $path -correct useGrid=false genomeSize=$genomesize minInputCoverage=1 \
-nanopore $raw_read
```

- Daccord v0.0.10

```
seqkit seq -w 150 $raw_read > $raw_read_split
fasta2DAM $reads.dam $raw_read_split
DBsplit -s256 -x1000 $reads.dam
HPC.daligner $reads.dam -t 24 > $reads.las
daccord $reads.las $reads.dam > $corrected_read
```

- CONSENT v2.2.2

```
CONSENT-correct --in $raw_read --out $corrected_read --type ONT
```
